# MinD proteins regulate CetZ1 localisation in *Haloferax volcanii*

**DOI:** 10.1101/2024.08.01.606189

**Authors:** Hannah J. Brown, Iain G. Duggin

## Abstract

CetZ proteins are archaea-specific homologues of the cytoskeletal proteins FtsZ and tubulin. In the pleomorphic archaeon *Haloferax volcanii*, CetZ1 contributes to the development of rod shape and motility, and has been implicated in the proper assembly and positioning of the archaellum and chemotaxis motility proteins. CetZ1 shows complex subcellular localization, including irregular midcell structures and filaments along the long axis of developing rods and patches at the cell poles of the motile rod cell type. The polar localizations of archaellum and chemotaxis proteins are also influenced by MinD4, the only previously characterized archaeal member of the MinD family of ATPases, which are better known for their roles in the positioning of the division ring in bacteria. Using *minD* mutant strains and CetZ1 subcellular localization studies, we show here that a second *minD* homolog, *minD2*, has a strong influence on motility and the localization of CetZ1. Knockout of the *minD2* gene altered the distribution of a fluorescent CetZ1-mTq2 fusion protein in a broad midcell zone and along the edges of rod cells, and inhibited the localization of CetZ1-mTq2 at the cell poles. MinD4 had a similar but weaker influence on motility and CetZ1-mTq2 localization. The MinD2/4 mutant strains formed rod cell shapes like the wildtype at an early log stage of growth. Our results are consistent with distinct roles for CetZ1 in rod shape formation and at the poles of mature rods, that are positioned through the action of the MinD proteins and contribute to the development of swimming motility in multiple ways. They represent the first report of MinD proteins controlling the positioning of tubulin superfamily proteins in archaea.

## Introduction

Spatial and temporal regulation of protein subcellular localization is critical for the proper function of all cells. In bacteria, there are several different classes of proteins from the ParA/MinD superfamily [1], which coordinate positioning of flagella [2; 3], chemosensory arrays [4; 5], chromosomes [6], plasmids [7], cell division cytoskeletal proteins [8; 9], and various other molecules or structures [10; 11; 12; 13; 14]. A key characteristic of ParA/MinD superfamily proteins is their ATPase activity [15; 16] and dimerization [9; 17] involving amino acid residues within a deviant Walker A motif [1; 10]. In MinD proteins, a subset of proteins within the ParA/MinD superfamily, this ATP binding and hydrolysis drives intracellular Min system dynamics that can generate concentration gradients of Min proteins [7; 15; 16]. In bacteria, these gradients are critical for the correct positioning of FtsZ, a key cell division protein and tubulin superfamily homologue [18; 19; 20], at mid-cell to help ensure symmetric division of the mother cell into two daughter cells [9; 21; 22].

The Min system of *Escherichia coli* is the prototypical example of Min system dynamics and function in bacteria. MinD oscillates between poles of the rod-shaped cells, controlled by its regulator MinE, and carries MinC an FtsZ antagonist. Together, these proteins generate dynamic concentration gradients of MinCD which is on average highest at cell poles and lowest at mid-cell, preventing Z-ring formation anywhere other than at mid-cell [8]. MinE accelerates the ATPase activity of MinD [15] and drives its release from the membrane which it is bound to via a membrane targeting sequence [23; 24; 25]. Together these proteins constitute a reaction-diffusion system driven by the hydrolysis of ATP. Disruption of MinD ATPase activity blocks oscillation and function completely [26; 27].

ParA/MinD family homologues have been identified across the archaeal domain, but MinD homologues are particularly abundant in *Euryarchaea* [28]. In the model halophilic archaeon, *Haloferax volcanii*, there are four MinD paralogues. The functions of MinD1-3 (HVO_0225, HVO_0595, and HVO_1634, respectively) have not been described and no clear protein partners equivalent to bacterial MinE have been identified. MinD4 (HVO_0322) has been characterised and is known to oscillate between cell poles, as well as concentrating with a cap-like appearance at the poles [28]. Surprisingly, the oscillation was not dependent on the ATPase Walker A or Walker B motifs, but the cap-like structures were, suggesting that the oscillation is driven by an alternative energy source [28]. Furthermore, although *H. volcanii* possess FtsZ homologues, none of the MinD paralogues were found to have a role in the mid-cell positioning of FtsZ; simultaneous deletion of all four paralogues had no detected effect of FtsZ positioning. Instead, deletion of *minD4* caused a motility defect and reduced the polar positioning of key protein constituents of the archaellum and chemotaxis arrays, ArlD and CheW, respectively [28]. Furthermore, disruption of MinD4 predicted ATPase activity through mutations to the Walker A or Walker B motifs, or truncation of its extended C-terminal tail all result in motility defects. Potential roles of the other MinD paralogues in motility of *H. volcanii* have not been reported.

Although the archaellum and chemosensory arrays are considered the core motility machinery in *H. volcanii*, there are many other contributing factors, and a range of proteins with a variety of primary functions have been shown to influence motility [29; 30; 31; 32; 33; 34; 35; 36; 37; 38]. CetZ cytoskeletal proteins, a group of tubulin superfamily proteins specific to archaea and abundant in Haloarchaea [37; 39; 40], have been implicated in the control of cell shape and motility [37; 41]. The most well-studied and highly conserved CetZ homologue, CetZ1, is necessary for rod-shape development and swimming motility [37; 41]. Our recent results have suggested that CetZ1 is an important factor for the proper assembly and positioning of motility proteins ArlD and CheW [42]. It is not yet clear whether CetZ1 contributes to motility solely via its essential role in rod shape development [37], or whether it has other functions that promote motility. It also is not yet clear how MinD4 and CetZ1 might coordinate their activities to regulate positioning and assembly of motility machinery.

Currently, no connection between Min proteins and FtsZ or other tubulin superfamily homologues, such as the CetZ, has been reported in archaea. Here, we investigate the potential for MinD homologues of *H. volcanii* to control positioning of CetZ1 and identify MinD2 as a strong regulator of CetZ1 localisation in rod cells. This is consistent with a model in which a hierarchy or network of proteins is involved the promoting the development of motility structures at the cell poles.

## Methods

### Growth and culturing

*Haloferax volcanii* was routinely grown in Hv-Cab [43] medium supplemented with uracil (50 μg/mL) to fulfil auxotrophic requirements if necessary, or L-Tryptophan (1 mM) to induce expression of genes under the control of the p.*tna* promotor. Liquid cultures were grown at 42 °C with shaking (200 rpm).

### Strain and plasmid construction

Strains, plasmids and oligonucleotides used in this study are listed in Tables S1, S2 and S3, respectively. *H. volcanii* H26 (Δ*pyrE2* ΔpHV2) was used as the wildtype parent strain for construction of mutant strains. All strains investigated in experiments contained a pHV2-based vector (pTA962 vector or derivatives) to complement the *pyrE2* auxotrophic marker and better match the original *H. volcanii* wild-type genotype; previous work has shown that the base strains with Δ*pyrE2*, Δ*hdrB* or ΔpHV2 show poorer expression of the cell morphology and motility phenotypes, which are sensitive to nutrient and culture conditions [37; 43; 44].

To create chromosomal point mutations in *minD2*, generating strains ID810 (*minD2WA\**) and ID811 (*minD2WB\**), H26 was transformed with vectors pTA131_minD2WA or pTA131_minD2WB, respectively, using previously described transformation methods [45] and the homologous recombination-based ‘pop-in pop-out’ method [46] was used to replace the native *minD2* gene with Walker A (K16A, AAG to GCC) or Walker B (D117A, GAT to GCC) mutant alleles. Primers MinD2_USflank_extF (forward) and MinD2_DSflank_extR (reverse), designed to bind externally to the regions of homologous recombination, were then used to amplify the *minD2* locus. The resultant PCR products were purified, and then sequenced by Sanger sequencing (Australian Genome Research Facility) to identify transformants carrying the expected mutations. All strains using *minD2WA\** (ID810) as the parent strain were also verified by checking for complete conversion of alleles and the absence of potential suppressor mutations (SNPs, indels) using Illumina whole-genome sequencing with ∼50X coverage.

Plasmids for the expression of *minD2, minD2WA\**, and *minD2WB\** were generated by amplifying these open reading frames from *H. volcanii* H26 (ID621), ID810, and ID811, respectively. Forward and reverse primers for this reaction incorporated NdeI and BamHI restriction sites at 5’ and 3’ ends, respectively. ORFs were then ligated between the same restriction sites in pTA962 [47], generating pHJB63-65 which were sequence verified.pHJB72 for the dual expression of *cetZ1-G-mTq2* and *minD2* was generated by amplifying the *minD2* open reading frame from pHJB63, incorporating an XbaI site at the 5’ end. The amplicon was then ligated between NheI and NotI of pHVID135. Plasmids and oligonucleotides used in this study are listed in Table S2 and S3, respectively.

### Soft-agar motility assays

Soft-agar motility assays were carried out as described in [42], using Hv-Cab with 0.25% (w/v) agar and 1 mM L-Tryptophan. Motility was quantified by averaging perpendicular measurements of the motility halo diameter after three days of incubation of the plates inside a plastic bag at 42 °C.

### Light microscopy

To prepare samples for microscopy, single colonies were first used to inoculate 3 mL Hv-Cab medium, each representing a single culture (biological) replicate. These cultures were allowed to grow to OD_600_ ∼0.6. Then, 3 μL of this culture was taken to inoculate 10 mL Hv-Cab supplemented with 1 mM L-Tryptophan and grown for 18 h (OD_600_ ∼ 0.1). A 2 μL droplet of cells was spotted onto a 1.5% agarose pad containing 18% BSW as described in [37]. Samples were then covered with a clean #1.5 glass coverslip.

Imaging for phase-contrast and fluorescence microscopy was carried out using a V3 DeltaVision Elite (GE Healthcare) with an Olympus UPLSAPO 100X NA 1.4 objective and pco.edge 5.5 sCMOS camera. A filter set for mTurquoise2/CFP (excitation=400-453, emission=463-487) was used for imaging of CetZ1-mTq2 expressing strains. 3D Structured-illumination microscopy (3D-SIM) was conducted as described previously [41] using a DeltaVision OMX SR microscope (GE Healthcare) and a DAPI filter for excitation (405 nm) and an AF488 filter for emission (504-552 nm), suitable for imaging of mTurquoise2. Raw images were reconstructed using SoftWorx (Applied Precision, GE Healthcare), and 3D renders were generated [48]using Bitplane IMARIS (v9.6.0).

### Image analysis

Image analysis was conducted using the MicrobeJ (v5.13I) [49] plugin for FIJI (v2.14.0/1.54f) [50]. Cell shape characteristics were analysed using phase-contrast images and default settings. Fluorescent foci detection was carried out after background subtraction with a ball size of 50 pixels, and the *point* detection method was used for analysis of CetZ1-mTq2 localisation (Tolerance=200).

Fluorescence intensity profiles along the long axis of each cell were generated using the *XStatProfile* function in MicrobeJ (the long axis was normalized between 0-1). The mean and standard deviation of the raw fluorescence intensity over all cells were then determined (Stat=mean±stdv, Y axis=MEDIAL.intensity.ch2, number of bins=50). The intensity data were also normalized (0-100%) and the median fluorescence intensity along the long axis determined for each culture replicate (Stat=median, Y axis=MEDIAL.intensity.ch2, number of bins=50), and then the mean and standard error of all culture replicates were plotted. These data were also displayed as the difference in fluorescence intensity between *minD* deletion or Walker mutant backgrounds and the wildtype background.

Heatmaps of foci localisation were generated using pooled data from all culture replicates and the *XYCellDensity* function (Rendering=Density) with default settings.

## Results

### Deletion of minD2 or minD4 results in motility defects with normal rod cell shape

We began by assessing the effect of *H. volcanii minD* gene deletions on motility, focussing on MinD2 and MinD4, which are expressed significantly in standard culture conditions [51; 52; 53]. MinD4 has previously been studied in this context [28]. The *minD* deletion strains grew at rates like the wildtype in starter cultures, and motility was assessed using soft-agar motility assays in which cells develop into a motile rod form [37] and then swim out through the gel from a central inoculation point, forming a halo.

Deletion of *minD2* resulted in a motility defect (approximately 25% of the halo diameter compared to wildtype), which was significantly stronger than that caused by deletion of *minD4* (approximately 50% of wildtype motility) (Fig. 1a, b). Deletion of both *minD2* and *minD4* resulted in a stronger motility defect (approximately 16% of wildtype motility). The deletion of all four *minD* homologues (Δ*minD1/2/3/4*) was indistinguishable to that of the Δ*minD2/4*.

**Figure 1.**
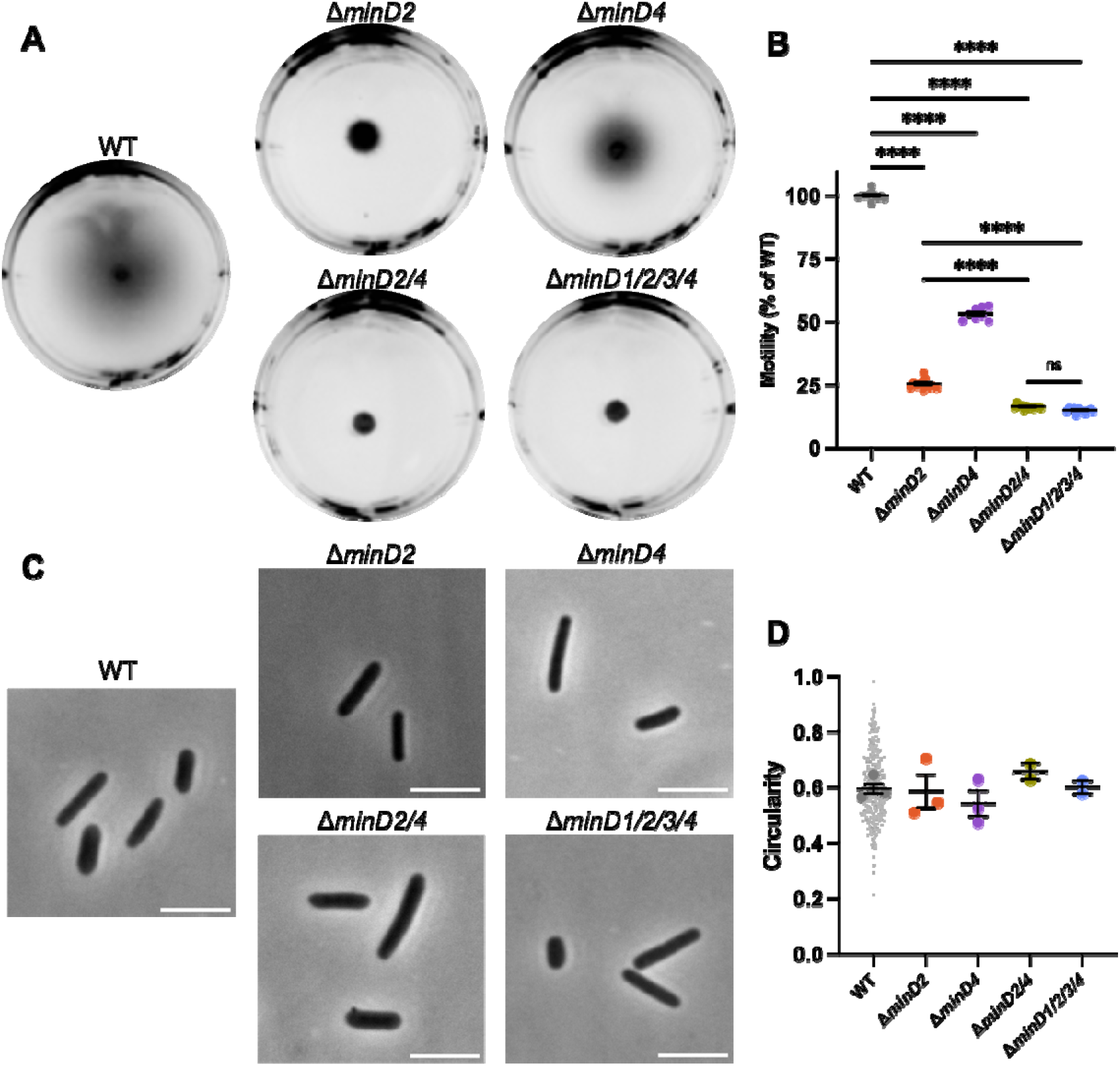
MinD2 and MinD4 have roles in motility without influencing shape. **A)** H26 wildtype and *minD* deletion mutant backgrounds, carrying an empty pTA962 vector, were used to inoculate the centre of a soft agar plate. **B)** The diameter of their motility halos was measured across six independent liquid starter culture replicates (represented by individual points), each determined from the mean of at least two replicate agar plates. Mean and standard error is shown. **C)** Phase-contrast microscopy of cells grown in Hv-Cab liquid medium with 1 mM L-tryptophan, sampled at OD_600_ ∼ 0.1. Scale bars = 5 μm. **D)** Cell circularity distributions, quantified from phase-contrast images as a proxy for cell elongation. Large data points represent mean circularity of one culture replicate, small data points represent individual cells from all culture replicates for the indicated strain. Mean and standard error of culture replicate means are shown (bars). The number of individual cells measured from pooled culture replicates were as follows: WT + pTA962, n=996; Δ*minD2* + pTA962, n=247; Δ*minD4* + pTA962, n=217; Δ*minD2/4* + pTA962, n=223; Δ*minD1/2/3/4* + pTA962, n=279. In **B** and **D**, one-way ANOVA was used as a statistical test, and only significant and/or relevant comparisons are shown.****p<0.00005, ns=not significant.

*H. volcanii* is pleomorphic and cultures can show a range of cell shapes including irregular disk, or plate-shaped cells, and rods with varying widths and degrees of elongation [37; 43]. Furthermore, a motile-rod cell type with regular cell widths develops under certain conditions to promote swimming [37; 54]. To determine whether the two MinD proteins are required for rod formation, which might contribute to their effects on motility, we sampled wild-type and Δ*minD2/4* cells from the early-log phase of liquid batch cultures (OD ∼ 0.1), where motile rods are prevalent [43; 54]. By quantifying cell circularity as an indicator of the extent of cell elongation, we observed no significant cell shape differences at this stage of growth in any of the strains (Fig. 1c, d).

### MinD2 and MinD4 influence the cellular positioning of CetZ1

MinD4 has been shown to have a role in the polar positioning of the archaellum and chemotaxis arrays through the use of tagged marker proteins ArlD and CheW, respectively [28]. We recently found that CetZ1 also influences the proper assembly and positioning of ArlD and CheW [42]. It is unknown whether MinD4 and CetZ1 cooperate or work independently to promote motility. To begin investigating this, we sought to determine whether the absence of MinD2 or MinD4 affects the localisation of CetZ1.

To do this, we observed the localisation of a CetZ1-mTurquoise2 (mTq2) fusion protein [55] in the wildtype and various *minD* deletion strain backgrounds. Fluorescence microscopy of CetZ1-mTq2 in wildtype cells imaged in early-log growth (OD_600_ ∼ 0.1) (Fig. 2a), showed the expected patchy localisation with relatively high fluorescence at cell poles and the mid-cell region [41; 42; 55]. To characterise these patterns, we analysed the whole population of cells, in the wildtype and *minD* mutant backgrounds, by generating heatmaps that represent the position and frequency of fluorescent foci within a normalized cell shape (reflecting the averaged aspect ratio) (Fig. 2b). We also generated plots of the total and normalized mean fluorescence intensity of CetZ1-mTq2 along the long-axis of the cells (Fig. 2c-d). In the Δ*minD2* background, CetZ1-mTq2 localisation remained patchy and concentrated in a broad zone near midcell, which was noticeably broader compared to the wildtype, and the high-intensity polar localisations of CetZ1-mTq2 were not observed (e.g., Fig. 2a, c, d). These patterns were also reflected in the number of fluorescent foci detected along the cell lengths (Fig. 2b); the wildtype background displayed clear peaks of fluorescence around midcell and at the cell poles, but in the Δ*minD2* background the midcell foci were more spread out along the cell length, and showed almost complete loss of the polar clusters of foci.

**Figure 2.**
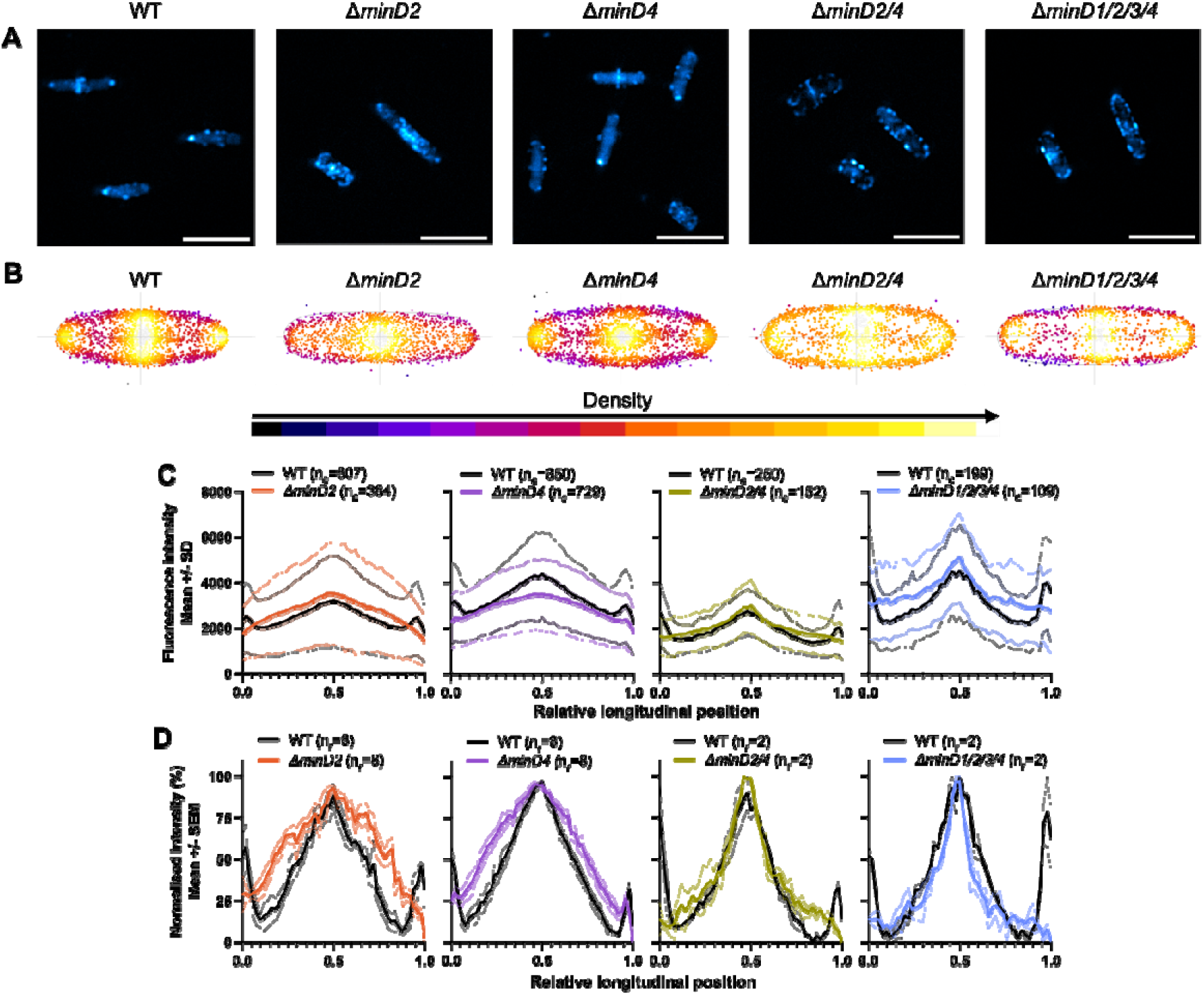
MinD2 controls polar positioning of CetZ1. H26 wildtype and minD deletion strains containing pHVID135 (for expression of CetZ1-mTq2) were grown in Hv-Cab with 1 mM L-Tryptophan and sampled at an approximate OD_600_ of 0.1. **A)** Epifluorescence images of CetZ1-mTq2 localisation in the indicated minD knockout strains (scale bar 5 μm). **B)** Heatmaps representing the position and number (heatmap density) of detected CetZ1-mTq2 fluorescence peak intensities (foci) along the long (vertical) and lateral (horizontal) axes of all cells combined. **C)** The median fluorescence intensity of CetZ1-mTq2 along the normalized long axis of each cell was used to generate a plot of the mean (bold line) and standard deviation (shading) of the raw fluorescence intensity data from all cells (n_c_) (combined from at least two replicate cultures). Cultures of the wildtype strain were repeated in parallel alongside each mutant independently. **D)** The same intensity data were also normalized (0-100%) for each culture replicate and plotted as the mean (bold line) and standard error (shading) of all culture replicates (n_r_) to enable comparison of protein distributions amongst the various strains.

To confirm whether the changes to CetZ1-mTq2 localization patterns were resultant from the loss of *minD2* specifically, we observed CetZ1-mTq2 localisation when *minD2* deletion was complemented by expression of *minD2* from a plasmid containing the open reading frames for *cetZ1-mTq2* and *minD2* in tandem, under the control of the inducible *p*.*tna* promotor (Fig. S1). Dual expression of these genes in the Δ*minD2* background using 1 mM L-Tryptophan did not clearly restore the CetZ1-mTq2 localisation defect (Fig. S1a). However, when 2 mM L-Tryptophan was used, the localisation of CetZ1-mTq2 was partially restored (Fig. S1c-g), confirming that *minD2* regulates the localization of CetZ1. The requirement for the additional tryptophan to observe partial complementation is likely due to a sensitivity of the phenotype to the expression level of *minD2*, which is likely to be influenced by its second position in the tandem expression plasmid [55].

A weaker defect of CetZ1-mTq2 polar localization was also observed in the Δ*minD4* background (compare the polar intensities between mutant and wildtype in Fig. 2d). Additional control experiments demonstrated that the Δ*minD* strains producing CetZ1-mTq2 had no significant differences in shape compared to the wildtype control (OD ∼ 0.1), and we also observed an equivalent motility defect with or without the CetZ1-mTq2 fusion (Fig. S2a, b).The results in Figure 2 also revealed that the zone of CetZ1-mTq2 localization around the midcell region was broadened in the absence of either MinD2 or MinD4 (Fig. 2b, d). Interestingly, such broadening was not apparent in strains in which both *minD2* and *minD4* were deleted, although the polar localization of CetZ1-mTq2 was still strongly inhibited (Fig. 2b, d). This suggests that the midcell zone of CetZ1-mTq2 localization is impacted when only one of MinD2 or MinD4 is present and implies a genetic interaction or interference between *minD2* and *minD4* that influences midcell CetZ1.

### Structural characteristics of CetZ1-mTq2 localization in individual cells lacking *minD2* or *minD4*

We further resolved structural characteristics of the CetZ1-mTq2 polar localizations and the effects of the *minD* knock-outs, using three-dimensional structured-illumination microscopy (3D-SIM). Multiple cells are shown as maximum intensity projections in Fig. 3a-c to visualize the effect of the mutations on CetZ1 structures in individual cells and facilitate comparisons to the aggregate data shown in Fig. 2b-d.

**Figure 3.**
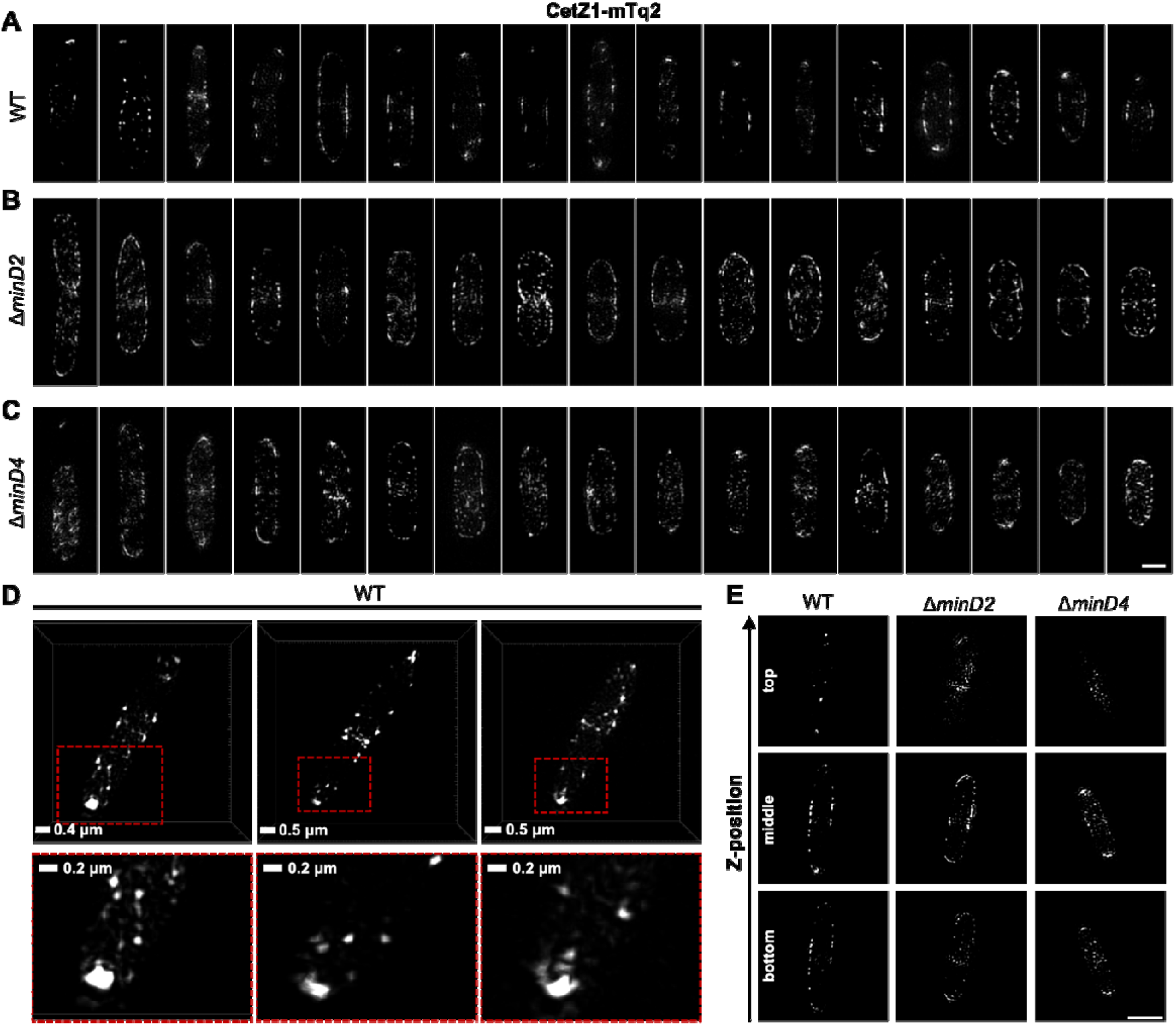
Subcellular structures of CetZ1-mTq2 in wildtype and *minD* mutant backgrounds, by 3D-SIM imaging. **(A-C)** Maximum intensity projections of CetZ1-mTq2 in H26 wildtype, Δ*minD2*, and Δ*minD4* backgrounds, viewed along the imaging z-axis (i.e., XY-views). Scale bar in panel C is 1 μm and applies to panels A-C. **D)** XY-views of 3D rendered images, showing non-circular fluorescence foci of CetZ1-mTq2 at the cell poles in H26 wildtype background. Red boxes indicate zoomed area (bottom). **E)** Examples of 3D-SIM slice sections of CetZ1-mTq2 in the indicated strains at approximate bottom, middle, and top z-positions of the cells (scale bar is 2 μm and applies to all images).

In wildtype early-log cells, CetZ1-mTq2 often appeared as a bright focus, doublet, multi-lobed, or cap-like structure at the cell poles, suggestive of a ring or disc-like structure at the tip of the cell pole (Fig. 3a, d). Patches and filaments were also observed along the cell edges in the midcell zone. In the absence of *minD2*, the foci of CetZ1-mTq2 appeared more randomly distributed around the cells, with no clear preference for the cell poles or the midcell zone(Fig. 3b). In the absence of *minD4*, this effect was weaker, and some of the cells still appeared to show the relatively bright polar tip structures of CetZ1-mTq2 (Fig. 3c). The Δ*minD2* and Δ*minD4* backgrounds also showed noticeably more of the CetZ1-mTq2 patches or foci throughout the cell area compared to the wildtype (Fig. 3a-c). By inspecting the slice series of images to resolve the 3D detail (Fig. 3e, Supplementary Video 4), CetZ1-mTq2 foci appeared noticeably more prevalent at the top and bottom surfaces along the length of the *minD* mutant cells, compared to the wildtype, and potentially in the cytoplasm too. These observations are consistent with the broadening of the midcell zone and reduced polar positioning of CetZ1-mTq2 in the *minD* mutants seen in the aggregated epifluorescence data (Fig. 2b-d).

### ATPase active-site residues of MinD2 are required for normal swimming motility and polar localization of CetZ1

MinD proteins typically rely on ATP hydrolysis to drive their dynamic localization patterns, such as oscillation between cell poles, generating concentration gradients which form the basis for their spatial and temporal control of downstream proteins [10; 27; 28]. When the Walker A and Walker B motifs of *H. volcanii* MinD4 were previously mutated to disrupt ATP binding and hydrolysis, swimming motility was reduced and polar localisation of MinD4 was disrupted, although not its oscillation [28].

Given that MinD2 had the strongest effects on swimming motility and CetZ1 localization, we chose to further investigate the importance of the ATPase active-site for MinD2 functionality in motility and CetZ1 localization. We generated chromosomal point-mutations in the endogenous *minD2* gene in the ATPase active site via the deviant Walker A (K16A) and Walker B (D117A) motifs, here forth referred to as *minD2WA\** and *minD2WB\**, respectively. These mutations are equivalent to those previously made in *minD4* [28].

Both *minD2WA\** and *minD2WB\** strains (containing the empty vector pTA962) partially perturbed motility, resulting in ∼78% and ∼36% of the wildtype motility halo diameter, respectively (Fig. 4a, c), compared to the complete deletion of *minD2* (∼25%, Fig. 1a, b). The motility defect of Δ*minD2* was complemented by expressing *minD2* from a plasmid (pTA962 backbone), confirming the important role of *minD2* in motility (Fig. 4b, c).

**Figure 4.**
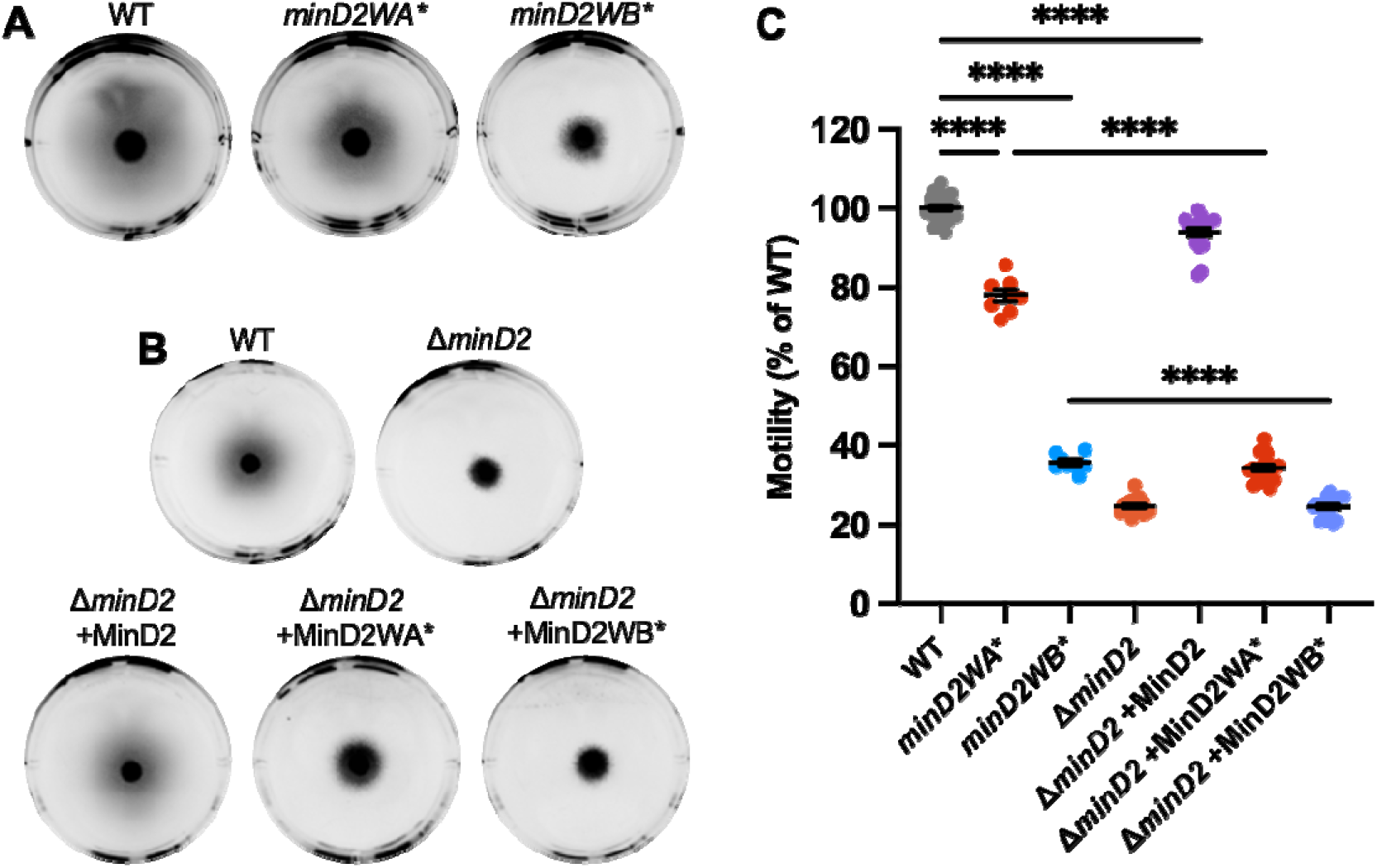
Chromosomal Walker mutants of *minD2* perturb motility. **A)** H26 wildtype, *minD2WA\**, and *minD2WB\** backgrounds containing pTA962, **B)** and Δ*minD2* containing pTA962 were compared to Δ*minD2* containing pHJB63-65 for expression of *minD2, minD2WA\**, and *minD2WB\**, respectively, and inoculated onto Hv-Cab soft agar (0.25 % w/v) supplemented with 1 mM L-Tryptophan. **C)** Halo diameters were quantified across four starter culture replicates with two technical replicates (motility agar plates) each (indicated by individual points). Mean and standard error are shown. One-way ANOVA was used as a statistical test, and only significant and relevant comparisons are shown. ****p<0.00005.

When *minD2WA*\* was expressed from a plasmid in the Δ*minD2* background, this strain had a more pronounced motility defect (∼35%) compared to the chromosomal mutant *minD2WA\** (∼78%). This might be related to different expression levels of *minD2WA\** in the plasmid with p.*tna* compared to its native chromosomal site. Expression of *minD2WB\** from pTA962 in Δ*minD2* produced equivalent motility to the complete *minD2* deletion, consistent with its much stronger effects on motility observed with the chromosomal *minD2WB*\* mutant (Fig. 4b, c).

Phase-contrast microscopy and analysis of cell shape (circularity) confirmed that, like Δ*minD2*, the *minD2WA\** and *minD2WB\** mutations did not significantly affect rod shape (at OD ∼0.1), whether expressed chromosomally (Fig. S3a, c) or from the plasmid (Fig. S3b, d).

We then assessed the effect of the *minD2WA\** and *minD2WB\** chromosomal mutants on CetZ1-mTq2 localisation. Control experiments showed that expression of CetZ1-mTq2 did not impact cell circularity or motility of either the *minD2WA\** or *minD2WB\** strains (Fig. S3c, e). Fluorescence microscopy (Fig. 5a) showed that both Walker mutations resulted in a substantial loss of CetZ1-mTq2 fluorescence intensity (Fig. 5c, dc) and foci localisation at cell poles and showed the midcell zone broadening (Fig. 5b) like in the Δ*minD2* background (Fig. 2).

**Figure 5.**
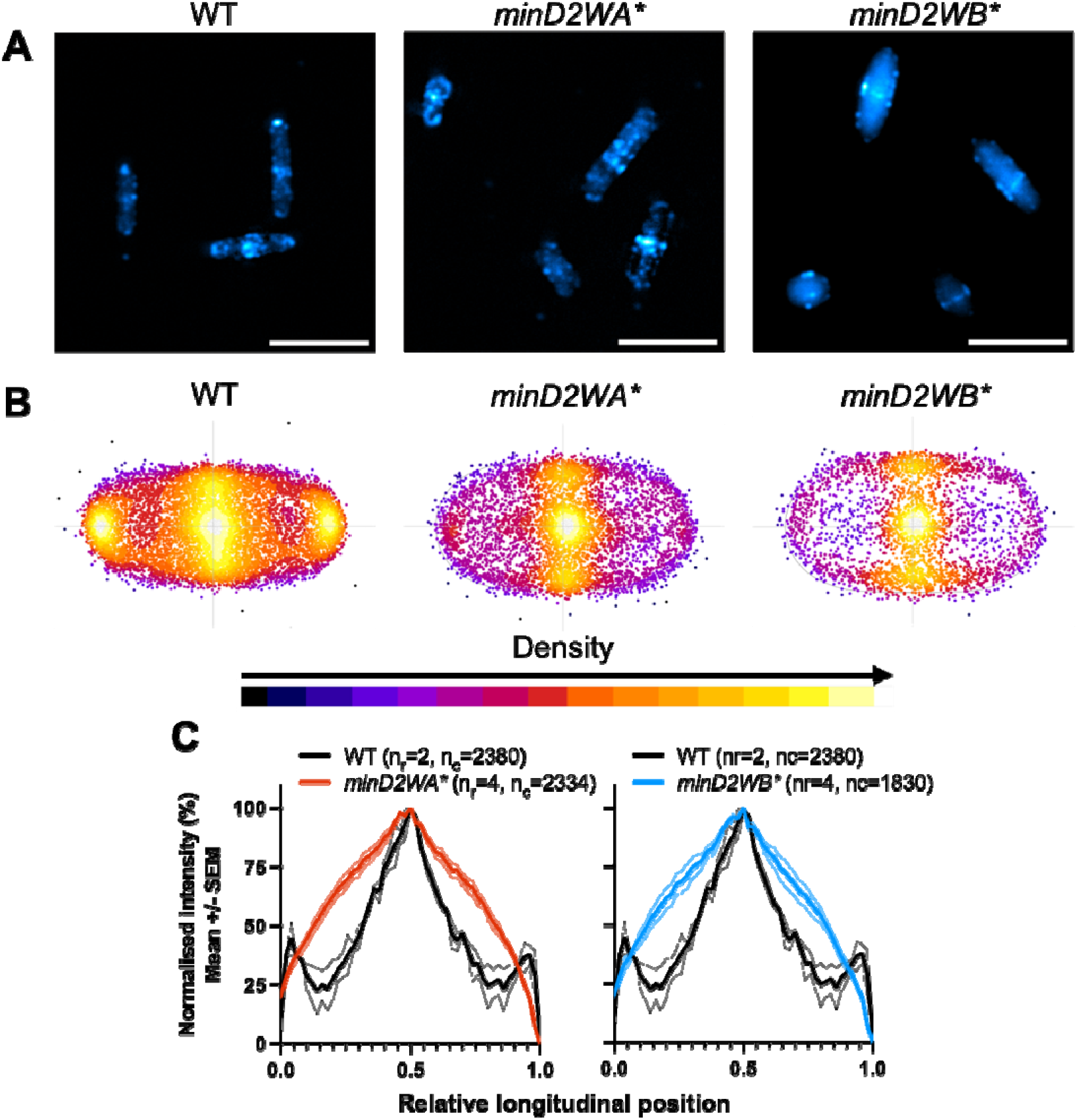
Chromosomal Walker mutants of *minD2* disrupt polar localisation of CetZ1. H26 wildtype and *minD2* walker mutant strains containing pHVID135 (for expression of CetZ1-mTq2) were grown in Hv-Cab with 1 mM L-Tryptophan and sampled at OD_600_ ∼ 0.1. **A)** Cells were imaged using epifluorescence to observe CetZ1-mTq2 localisation. Scale bar 5 μm. **B)** Heat map representations of detected foci of CetZ1-mTq2 and their localisations. Heatmaps are coloured by density, referring to subcellular localisation along the relative length of lateral and longitudinal axes only and do not fluorescence intensity. **C)** normalised median fluorescence intensity of CetZ1-mTq2 along the long axis of the cell, represented as the mean (bold line) and standard deviation (shading) of biological replicates (as in Fig. 2d). Two biological replicates were carried out for the wildtype background, and four for each of *minD2WA\** and *minD2WB\**, and data for H26 wildtype is included on both graphs for reference. The number of cells measured from pooled biological replicates is as follows: WT + CetZ1-mTq2, n=2380; *minD2WA\** + CetZ1-mTq2, n=2334; *minD2WB\** + CetZ1-mTq2, n=1830.

## Discussion

MinD proteins and related protein families are important for spatial and temporal positioning of a variety of molecules but have mainly been characterized in bacteria [10]. One of four MinD paralogues in the archaeal model organism *H. volcanii*, MinD4, has previously been described to affect motility via regulating the positioning of archaellum and chemotaxis arrays, via unknown mechanisms [28]. The tubulin-like cytoskeletal protein CetZ1 is another factor promoting the proper assembly and positioning of these proteins at the cell poles [42]. CetZ1 is needed for rod cell shape formation [37], which may contribute to motility in several ways, and shows complex subcellular localization along the cell body and then at the cell poles of motile rods, where it is hypothesised to form a scaffold-like structure, though its mechanism of action at these sites is unclear.

We found here that MinD2 is important for motility and for the subcellular positioning of CetZ1. The potential relationship between these two functions is yet to be elucidated. Deletion of *minD2* was found to have a more severe motility defect than deletion of *minD4*, and deletion of both genes resulted in an even greater motility defect, suggesting distinct roles or additive contributions of these two MinD paralogues in motility. MinD2 and to a lesser extent MinD4 also influenced the subcellular positioning of CetZ1 in the midcell region and at the poles, in a manner that was independent of cell shape. CetZ1 has been implicated in distinct roles in establishing rod-shape during motile cell differentiation, and then at the poles of the mature motile rods [37; 42]. Our results suggest that the effects of MinD2 on CetZ1 localization do not have an essential role in maintaining rod shape, because Δ*minD2* cells could still form normal shaped rods at an early-mid log stage of culture.

We found that mutations in the Walker A and B motifs of the MinD2 ATPase both strongly influenced CetZ1 positioning along the cell length and poles, like the *minD2* deletion. However, only the Walker B mutant (D117A) strongly perturbed motility, whereas the Walker A (K16A) mutant (in its chromosomal context) resulted in only a mild motility defect. This suggests that MinD2 has an impact on motility that is largely independent of the K16 residue and CetZ1 positioning. The K16 residue is expected to be involved in nucleotide dependent dimerization [27], so this function may not be critical to the overall motility function of MinD2, and instead plays a role in CetZ1 positioning for an as yet unresolved purpose. Notably, the equivalent MinD4 Walker A and B mutants reduced motility equivalently to *minD4* deletion [28]. The MinD2 and MinD4 proteins could therefore have multiple, differing functions that contribute to the proper development of motile *H. volcanii* cells.

The accompanying study by Patro et al. [56] found that MinD2 is required for normal motility and the proper polar localisation of the archaellum and chemotaxis proteins, ArlD and CheW, like previously seen with MinD4 [28]. They also reported a difference in cell shape during the development of rod cells in the Δ*minD2* compared to the wildtype. This likely represents an earlier phase of *H. volcanii* cultures, where cells are actively shapeshifting into motile rods [43; 54], compared to the current study where rods had already formed, and we found that their cell shapes in the wildtype and Δ*minD2* strains were indistinguishable. Though we note that since *H. volcanii* morphogenesis is highly sensitive to exact media composition and culture conditions [43; 44], excessive comparisons of the exact magnitudes or timing of events from separate studies should be avoided. Nevertheless, if the rate or timing of rod formation is influenced by MinD2 [56], this may affect another developmental stage of motility, such as the placement of Arl/Che proteins at the poles. However, it is still unclear whether any effects on cell shape development directly contribute to the *minD2*-dependent reduction of motility.

Considering the main findings from this and other recent studies [28; 56], we can summarize several possibilities for the role of MinD proteins in motility and CetZ1 localization (Fig. 6). In this model, MinD2 and MinD4 help direct the archaellum (Arl) and chemotaxis (Che) proteins to the cell poles directly or via positioning and assembly of polar CetZ1. We recently found that CetZ1 plays a role in the proper placement of the motility machinery [42]. However, the present finding that the chromosomal *minD2WA\** mutation blocked polar CetZ1 localization but only weakly affected motility suggests that MinD proteins might separately control both the position of CetZ1 and another aspect of motility, such as placement of the motility machinery at the cell poles, independently of MinD2 regulated CetZ1 positioning.

**Figure 6.**
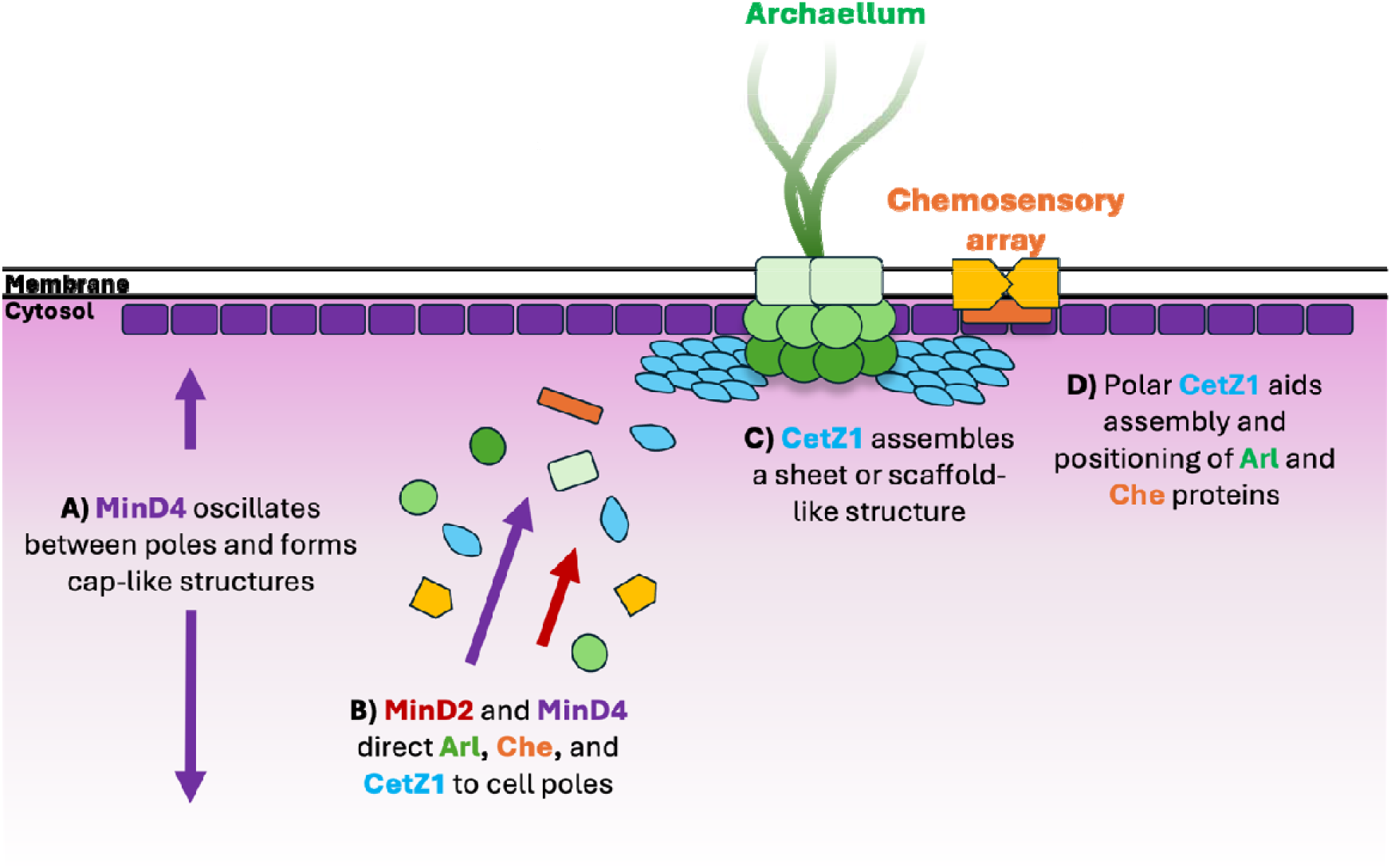
Schematic model for assembly of motility proteins at cell poles. **A)** MinD4 oscillates between cell poles and forms cap-like localisations at cell poles in an ATPase dependent manner [28]. **B)** MinD4 [28] and MinD2 direct protein constituents of the archaellum (“Arl”) and chemosensory arrays (“Che”) such as ArlD and CheW, respectively, to cell poles [56]. CetZ1 is also directed to cell poles, likely by both MinD paralogues, but more dominantly by MinD2. **C)** At cell poles, CetZ1 is hypothesised to assemble as a sheet or disc-like structure, **D)** where it may contribute to assembly and positioning of motility structures via an unknown mechanism [42].

Previous work showed that MinD4 forms dynamic cell cap-like structures in an ATPase-dependent manner, and yet oscillates in the cells independently of ATPase function (Fig. 6), suggesting that it interacts with another oscillator that drives the movement [28]. The localization patterns of MinD2 are currently unknown due to the absence of functional fluorescently tagged MinD2 [56]. However, our observations that the complex patterns of CetZ1 localization are affected by MinD2 at multiple sites and at a cellular scale would suggest that similar cellular level MinD2 pattern formation might exist.

In summary, our findings have showed that multiple MinD homologues can have differing functions and regulate the subcellular positioning of the archaeal tubulin-like protein CetZ1 in the context of the development of swimming motility. They represent the first demonstration of MinD proteins controlling the position of tubulin superfamily proteins in archaea. The MinD-controlled positioning of CetZ1 in distinct subcellular locations is consistent with multiple functions of CetZ1, evoking parallels with the multiple functions of tubulins and other eukaryotic cytoskeletal proteins that are extensively regulated in time and space.

## Acknowledgements

The authors would like to acknowledge facilities support from L. Cole and A. Bottomley and the UTS Microbial Imaging Facility. The work was supported by the Australian Research Council (DP160101076 and FT160100010 to IGD).

## Author Contributions

Conceptualisation – HJB & IGD; Data curation - HJB; Formal Analysis – HJB; Investigation – HJB; Methodology – HJB & IGD; Supervision – IGD; Original draft – HJB; Revision and reviewing the manuscript – HJB & IGD; Funding Acquisition – IGD.

## Notes

### Competing Interest Statement

The authors have declared no competing interest.

### Summary of Updates

Inclusion of complementation data for minD2 effects on CetZ1 localization. Clarification of results and discussion. Other minor modifications.

