## Supplementary Information for "MinD proteins regulate CetZ1 localisation in *Haloferax volcanii*"

**Contents:**

**Legends for Supplementary Videos 1-4**

**Supplementary Figures**

1. Polar localisation of CetZ1-mTq2 can be partially restored by complementation of minD2 deletion.
2. CetZ1-mTq2 has no effect on motility or rod-shape in *minD* deletion backgrounds.
3. Cell circularity and motility analysis of chromosomal and plasmid-expressed minD2 Walker mutants.

**Supplementary Tables**

1. Strains used in this study
2. Plasmids used in this study
3. Oligonucleotides used in this study

**Legends for Supplementary Videos:**

1. Z-stack of 3D-SIM reconstructions showing CetZ1-mTq2 localization in WT (top), Δ*minD2* (middle), and Δ*minD4* (bottom). The first frame is the bottom-most Z-position, and the last frame is the top-most Z-position.
2. 3D render of 3D-SIM images showing CetZ1-mTq2 localization in the WT genetic background.
3. 3D render of 3D-SIM images showing CetZ1-mTq2 localization in the Δ*minD2* genetic background.
4. 3D render of 3D-SIM images showing CetZ1-mTq2 localization in the Δ*minD4* genetic background.

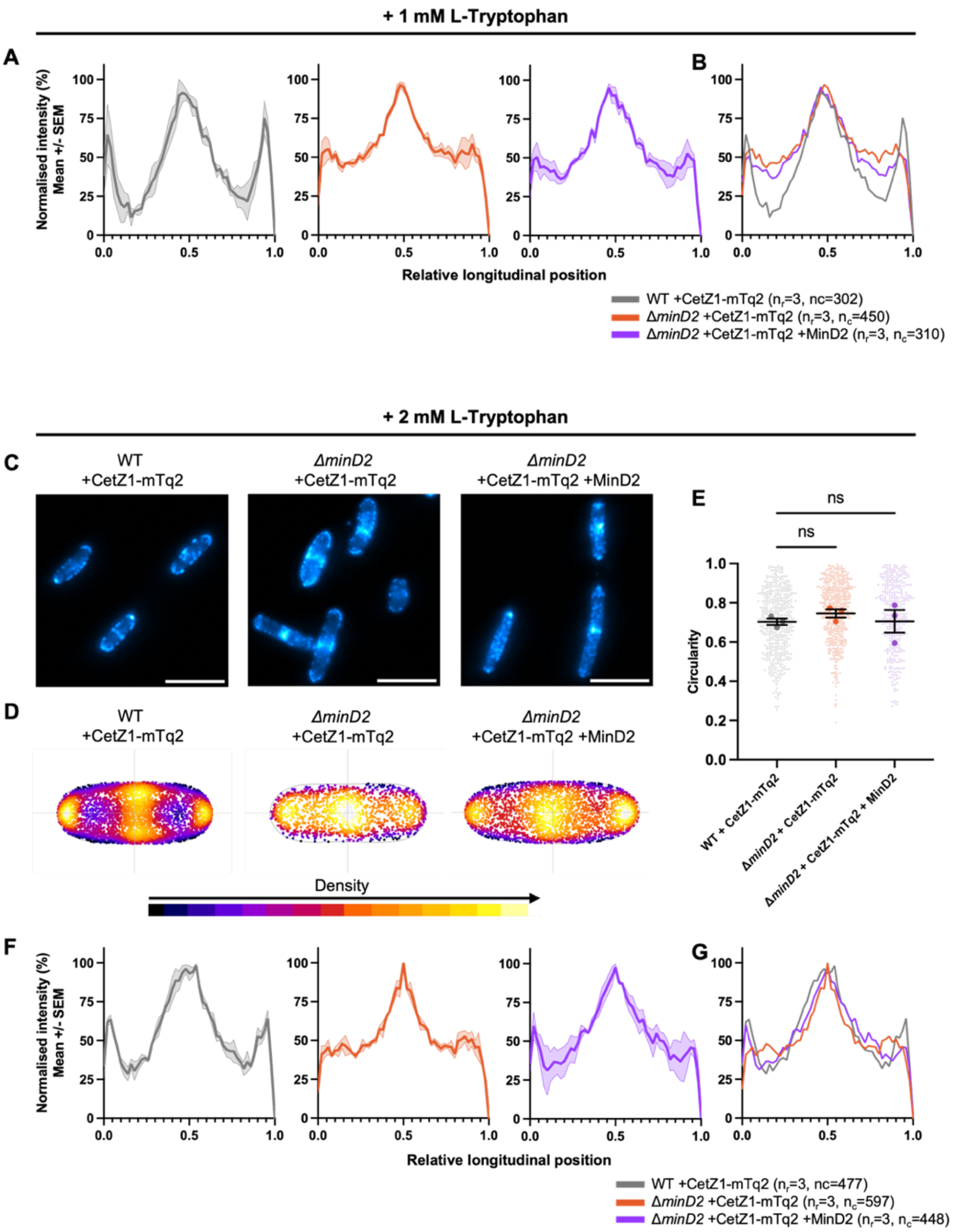


**Figure S1. Polar localisation of CetZ1-mTq2 can be partially restored by complementation of *minD2* deletion.** CetZ1-mTq2 was produced in the wildtype (H26) and Δ*minD2* backgrounds from pHVID135. MinD2 and CetZ1-mTq2 were simultaneously produced from pHJB72, containing the *cetZ1-mTq2* ORF immediately downstream from the *p.tna* promotor, followed by *minD2* in a tandem configuration. Three independent culture replicates ($n_c$) were carried out for each strain, and cells were sampled for imaging from Hv-Cab medium supplemented with 1 mM **(A-B)** or 2 mM **(C-F)** L-Tryptophan inducer, at an approximate $OD_{600}$ of 0.2. **A)** The median fluorescence intensity of CetZ1-mTq2 along the normalised long axis of each cell. Data were normalised (0-100%) for each culture replicate and plotted to show the mean (bold line) and standard error (shading) of all culture replicates. **B)** Overlay of the mean data in **A.** The number of culture replicates ($n_r$) and total number of individual cells analysed is indicated in the key. **C)** Fluorescence microscopy of CetZ1-mTq2 in the wildtype background, Δ*minD2* background, and Δ*minD2* background when co-produced with MinD2 for complementation. **D)** Heatmaps representing the position and number (heatmap density) of detected CetZ1-mTq2 fluorescence peak intensities (foci) along the long (vertical) and lateral (horizontal) axes of all cells combined. **E)** Superplots showing cell circularity measurements of each strain. Small individual points represent individual cells, large individual points represent mean cell circularity values of culture replicates. One-way ANOVA was used as a statistical test, ns=not significant. **F** Is as per **A**, and **G** as per **B,** but with 2 mM L-Tryptophan.

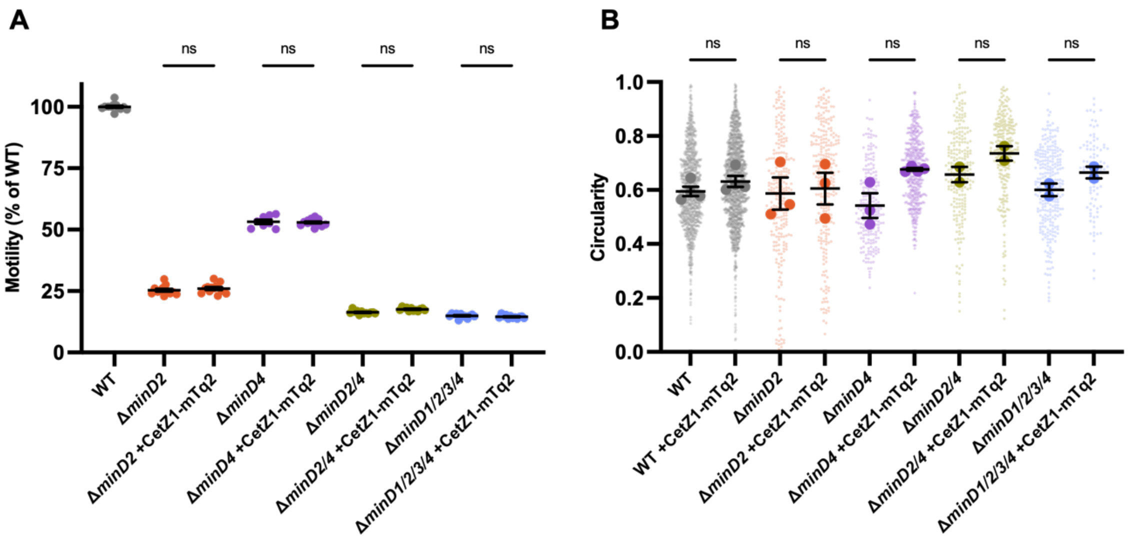


**Figure S2. CetZ1-mTq2 has no effect on motility or rod-shape in *minD* deletion backgrounds.** Wildtype and *minD* deletion strains containing the empty vector pTA962 or pHVID135 for expression of CetZ1-mTq2 were subjected to soft-agar motility assays and cell shape quantification. The relevant data from Fig. 1 and 4 are included here for reference. **A)** Quantification of halo diameter from Hv-Cab + 1 mM L-Tryptophan soft-agar (0.25%) motility assays. Individual points represent biological replicates. Mean and standard error are shown. **B)** Super plot showing cell circularity of cells grown in Hv-Cab medium with 1 mM L-Tryptophan and sampled at an approximate $OD_{600}$ of 0.2. Large data points represent mean circularity of one culture replicate, small data points represent individual cells from all culture replicates for the indicated strain. Mean and standard error of culture replicate means is shown. The number of individual cells measured from pooled biological replicates for *minD* deletion strains containing pTA962 are detailed in the legend of Fig. 1, and for remaining strains are as follows: **(B)** WT + CetZ1-mTq2, n=1341; Δ*minD2* + CetZ1-mTq2, n=271; Δ*minD4* + CetZ1-mTq2, n=432; Δ*minD2/4* + CetZ1-mTq2, n=251; Δ*minD1/2/3/4* + CetZ1-mTq2, n=97. One-way ANOVA was used as a statistical test (n=3), ns=not significant.

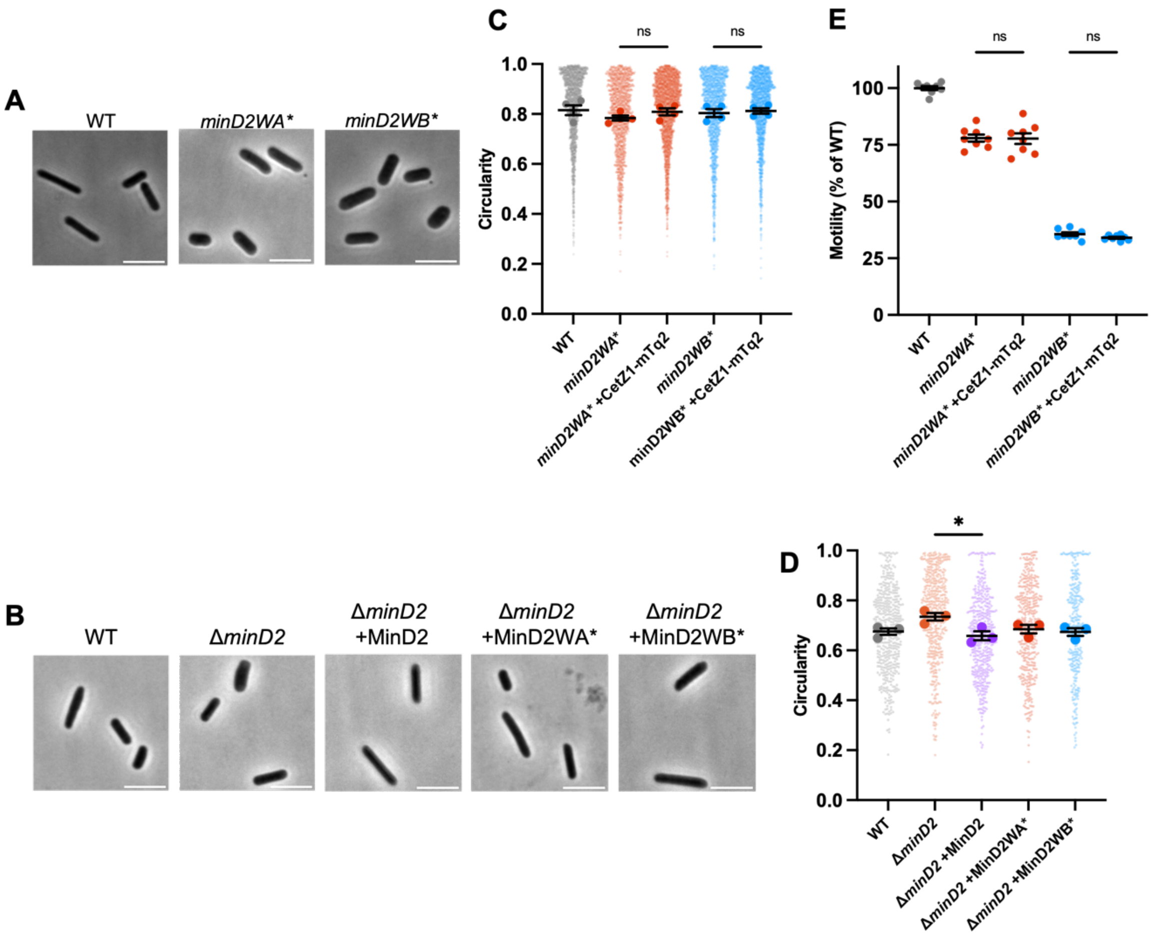


**Figure S3. Cell circularity and motility analysis of chromosomal and plasmid-expressed *minD2* Walker mutants. A)** Chromosomal *minD2* Walker mutants containing pTA962 and **B)** Δ*minD2* containing pHJB63-65 for expression of *minD2*, *minD2WA**, and *minD2WB**, respectively, were grown to an approximate $OD_{600}$ of 0.2 in HvCab medium supplemented with 1 mM L-Tryptophan. Representative phase-contrast microscopy images of *minD2* chromosomal walker mutants (scale bar 5 μm). **C)** Super plot showing cell circularity quantification of cells in **A** and *minD2* chromosomal Walker mutants with CetZ1-mTq2 (from Fig. 5a). Large datapoints represent mean circularity of all cells within one culture replicate. Small data points represent individual cells from all culture replicates in the indicated strain. Mean and standard deviation of culture replicate means is shown. The total number of cells analysed from pooled biological replicates is as follows: WT + pTA962, n=1851; minD2WA* + pTA962, n=1488; minD2WA* + CetZ1-mTq2, n=2334; minD2WB* + pTA962, n=1649; minD2WB* + CetZ1-mTq2, n=1830; **D)** As in **C**, but for strains shown in **B.** The total number of cells analysed from pooled biological replicates is as follows: WT + pTA962, n=446; Δ*minD2* + pTA962, n=500; Δ*minD2* +MinD2, n=456; Δ*minD2* +MinD2WA*, n=428; Δ*minD2* +MinD2WB*, n=319. **E)** Quantification of halo diameter from Hv-Cab + 1 mM L-Tryptophan soft-agar (0.25%) motility assays for strains in **A/C.** Individual points represent biological replicates. Data from Figure 4c is included here for reference. Mean and standard error are shown. In **C-E,** one-way ANOVA was used as a statistical test (n=3 for **C** and **D**). Only relevant comparisons are shown. *p<0.05, ns=not significant.

**Table S1.** Strains used in this study

| Strain | Description | Source |
|---|---|---|
| ID621 (H26) | Wildtype strain, cured of pHV2 and *ΔpyrE2* | [1] |
| HTQ228 (Δ*minD2*) | H26 with deletion of *minD2* (HVO_0595) | [2] |
| HTQ218 (Δ*minD4*) | H26 with deletion of *minD4* (HVO_0322) | [2] |
| HTQ241 (Δ*minD2/4*) | H26 with deletion of *minD2* (HVO_0595) and *minD4* (HVO_0322) | [2] |
| HTQ262 (Δ*minD1/2/3/4*) | H26 with deletion of *minD1* (HVO_0225), *minD2* (HVO_0595), *minD3* (HVO_1634), and *minD4* (HVO_0322) | [2] |
| ID810 (*minD2WA**) | H26 with a point mutation (K16A) within the walker A motif of *minD2* | This study |
| ID811 (*minD2WB**) | H26 with a point mutation (D117A) within the walker B motif of *minD2* | This study |

**Table S2.** Plasmids used in this study

| Plasmid | Description | Source |
|---|---|---|
| pTA962 | Expression vector containing *p.tna* promotor for *H. volcanii* and an *E. coli* shuttle plasmid | [3] |
| pHVID135 | For expression of CetZ1-G-mTurquoise2 fusion protein | [4] |
| pTA131_minD2WA | pTA131-based vector containing flanking sequences and Walker A mutation for minD2. Used to generate the chromosomal Walker A mutant minD2 strain *minD2WA**. | This study, Genscript |
| pTA131_minD2WB | pTA131-based vector containing flanking sequences and Walker B mutation for minD2. Used to generate the chromosomal Walker B mutant minD2 strain *minD2WB**. | This study, Genscript |
| pHJB63 | For expression of *minD2* from pTA962 (between NdeI and BamHI). | This study |
| pHJB64 | For expression of *minD2WA** from pTA962 (between NdeI and BamHI). | This study |
| pHJB65 | For expression of *minD2WB** from pTA962 (between NdeI and BamHI). | This study |
| pHJB72 | For dual expression of *cetZ1-mTq2* and *minD2* under the control of the same *p.tna* promotor. The ORF for *minD2* is cloned between NheI and NotI restriction sites of pHVID135, putting it in the second position behind *cetZ1-mTq2*. | This study |

**Table S3.** Oligonucleotides used in this study

| Name | Sequence (5'-3') | Use |
|---|---|---|
| MinD2_USflank_extF | CGACGTCGATTGCGTCCT | For amplification of *minD2* to confirm chromosomal point mutation of Walker A and Walker B motifs. |
| MinD2_DSflank_extR | GCTTTTGAACCGCTCTGCGA | For amplification of *minD2* to confirm chromosomal point mutation of Walker A and Walker B motifs. |
| NdeI-MinD2_F | GGCGGCCATATGGTCGAGGCGTTCGCC | For amplification of *minD2* to incorporate an NdeI site at the 5' end of the product. Used to generate pHJB63-65. |
| MinD2_BamHI_R | GGCGGCGGATCCTCATATTCGCTCGGGGACG | For amplification of *minD2* to incorporate a BamHI site at the 3' end of the product. Used to generate pHJB63-65. |
| XbaI_MinD2_F | GGCGGCTCTAGAATGGTCGAGGCGTTCGCC | For amplification of *minD2* to incorporate a XbaI site at the 5' end of the product. Used to generate pHJB72. |
| T3 | GCAATTAACCCTCACTAAAGG | For amplification of *minD2* for the generation of pHJB72. |
